# Impaired cytotoxic CD8+ T cell response in elderly COVID-19 patients

**DOI:** 10.1101/2020.08.21.262329

**Authors:** Jaana Westmeier, Krystallenia Paniskaki, Zehra Karaköse, Tanja Werner, Kathrin Sutter, Sebastian Dolff, Marvin Overbeck, Andreas Limmer, Jia Liu, Xin Zheng, Thorsten Brenner, Marc M. Berger, Oliver Witzke, Mirko Trilling, Mengji Lu, Dongliang Yang, Nina Babel, Timm Westhoff, Ulf Dittmer, Gennadiy Zelinskyy

**Affiliations:** Institute for Virology, University Hospital Essen, University of Duisburg-Essen, Essen, Germany; Center for Translational Medicine, Medical Department I, Marien Hospital Herne, University Hospital of the Ruhr-University Bochum, Herne, Germany; Department of Infectious Diseases, West German Centre of Infectious Diseases, University Hospital Essen, University Duisburg-Essen, Germany; Department of Anesthesiology, University Hospital Essen, University Duisburg-Essen, Germany; Department of Infectious Diseases, Union Hospital of Tonji Medical College, Huazhong University of Science and Technology, Wuhan, China; Joint International Laboratory of Infection and Immunity, HUST, Wuhan, China; Charité - Universitätsmedizin Berlin, Corporate Member of Freie Universität Berlin, Humboldt-Universität zu Berlin, and Berlin Institute of Health, Berlin-Brandenburg Center for Regenerative Therapies, Berlin, Germany; Medical Department I, Marien Hospital Herne, University Hospital of the Ruhr University of Bochum, Bochum

**Author notes:** Corresponding authors: Gennadiy Zelinskyy and Ulf Dittmer, Institute for Virology, University Hospital Essen, University of Duisburg-Essen, Essen, Germany.

**Keywords:** COVID-19, SARS-CoV-2, cytotoxic T cells, CD8+, CD4+, PD-1, granzyme, perforin, aging

## Abstract

SARS-CoV-2 infection induces a T cell response that most likely contributes to virus control in COVID-19 patients, but may also induce immunopathology. Until now, the cytotoxic T cell response has not been very well characterized in COVID-19 patients.

Here, we analyzed the differentiation and cytotoxic profile of T cells in 30 cases of mild COVID-19 during acute infection. SARS-CoV-2 infection induced a cytotoxic response of CD8+ T cells, but not CD4+ T cells, characterized by the simultaneous production of granzyme A and B, as well as perforin within different effector CD8+ T cell subsets. PD-1 expressing CD8+ T cells also produced cytotoxic molecules during acute infection indicating that they were not functionally exhausted. However, in COVID-19 patients over the age of 80 years the cytotoxic T cell potential was diminished, especially in effector memory and terminally differentiated effector CD8+ cells, showing that elderly patients have impaired cellular immunity against SARS-CoV-2.

Our data provides valuable information about T cell responses in COVID-19 patients that may also have important implications for vaccine development.

**Importance:** Cytotoxic T cells are responsible for the elimination of infected cells and are key players for the control of viruses. CD8+ T cells with an effector phenotype express cytotoxic molecules and are able to perform target cell killing. COVID-19 patients with a mild disease course were analyzed for the differentiation status and cytotoxic profile of CD8+ T cells. SARS-CoV-2 infection induced a vigorous cytotoxic CD8+ T cell response. However, this cytotoxic profile of T cells was not detected in COVID-19 patients over the age of 80 years. Thus, the absence of a cytotoxic response in elderly patients might be a possible reason for the more frequent severity of COVID-19 in this age group in comparison to younger patients.

## Introduction

Severe acute respiratory syndrome coronavirus 2 (SARS-CoV-2) is a highly virulent sarbecovirus currently causing a global pandemic with millions of cases and hundred thousands of fatalities. Virus replication in the lung epithelium and the corresponding pneumonia are the main reasons for symptomatic COVID-19 cases, although other tissues and organs such as the kidney are also affected (1). Elderly people are predisposed to severe COVID-19 and the mortality increases dramatically with age (1–3) . In particular, individuals over 80 years of age show the highest hazard ratio (8.93-13.77) in terms of hospital admissions (1) and have the highest case fatality rate (4).

There is growing evidence that adaptive immune responses are necessary for the control and subsequent elimination of the virus (5). Cytotoxic T lymphocytes (CTL) are a specialized population of immune cells which is able to selectively kill infected cells and consequently eliminate viruses. Usually, CD8+ T lymphocytes mediate adaptive cytotoxic T cell responses. Additionally, a fraction of the CD4+ T cell population is able to differentiate into cells with cytotoxic properties (6). Both populations of cytotoxic cells can contribute to virus control by eliminating infected cells. T cells responding to viral antigens expand and differentiate from cells with a naïve phenotype into subpopulations of terminally differentiated cytotoxic effector T cells or cells with an effector memory phenotype. Both effector cell subpopulations are abundant during the acute phase of antiviral immune responses (7). Accordingly, the number of cells with these phenotypes rises during the acute immune responses against several respiratory viral infections (8). The SARS-CoV-2 infection is associated with a reduction of CD8+ and CD4+ T cells (9, 10). One prominent cause of lymphopenia may be an enhanced migration of T cells into infected compartments (11, 12). Despite the lymphopenia, expanded virus-specific CD8+ and CD4+ T cells can be detected in COVID-19 patients (13, 14). The CD4+ and CD8+ T cells are specific towards several proteins of SARS-CoV-2 as has been recently shown (15–17). During the early phase of the immune response, CD8+ and CD4+ T cells reacted against the spike, membrane, and nucleocapsid proteins (15, 16). The T lymphocytes of convalescent patients responded to structural proteins or nonstructural proteins which provides evidence of the development of memory to different viral proteins after infection (17, 18). Interestingly, some individuals who were not infected with SARS-CoV-2 also responded to the antigens of this virus which have a low homology with “common cold” human coronaviruses (17, 18).

The detection of these virus-specific cells was possible after the *in vitro* stimulation of T cells with viral peptides. This method allows for the definition of the specificity of analyzed T cells, but has a modulating impact on the T cell phenotype and functionality. Moreover, the stimulation of activated effector T cells *in vitro* can lead to restimulation-induced cell death (RICD) (19). In our study, we have characterized lymphocytes without any treatment and performed ex vivo multiparameter analyses of T cells.

A key mechanism of functional CTLs is the elimination of virus-infected cells through the induction of apoptosis of target cells after cell-to-cell contact with effector CD8+ T cells. To perform cytotoxic functions, CTLs produce and accumulate effector molecules like the serine proteases granzymes (Gzm) and the pore-forming protein perforin in cytotoxic granules. Additionally, the release of Gzms from activated T cells contributes to the development of inflammation in infected organs. Gzms also change the intracellular matrix and support the migration of lymphocytes, while perforin is necessary for the entry of Gzms into target cells. After the formation of an immunological synapse and the degranulation of cytotoxic granules, Gzms enter into target cells where they initiate multiple pathways leading to the cell death of the infected cell, terminating intracellular virus replication through the loss of the host cell. Thus, the expression of different cytotoxic molecules in T cell subpopulations is an important hallmark for the existence of lymphocytes with a cytotoxic potential known to be necessary for virus control (20). Besides their beneficial role, the elimination of virus-infected cells by cytotoxic T cells can also be associated with a damage of infected organs resulting in severe immunopathology. Therefore, they are tightly controlled by multiple checkpoints of the immune system (21). Accordingly, the contribution of T cells to the pathology seen in COVID-19 patients has been recently discussed (22). Immune senescence is an age-associated change of the immune system related to thymus involution and reduced frequencies of naïve CD8+ and CD4+ T cells (23). The senescent immune system exhibits a reduced plasticity and adaptive effector potential to respond against viral infections (12). In the current study, we characterized the differentiation status of T lymphocytes and their production of cytotoxic molecules in 30 COVID-19 patients with a mild disease course. CD8+ T cells, but not CD4+ T cells, developed a cytotoxic phenotype during early SARS-CoV-2 infection. We observed reduced frequencies of T cells producing cytotoxic molecules in elderly patients. The ability to simultaneously produce Gzms and perforin was significantly impaired in aged patients.

## Results

To characterize the cytotoxic profile of T cells upon SARS-CoV-2 infection, we analyzed the blood of 30 COVID-19 patients with a mild disease course by multi-parameter flow cytometry. All analyses were performed directly *ex vivo* from blood cells to determine the *in vivo* phenotype of T cells in COVID-19 patients excluding changes that are inevitably induced by re-stimulation protocols *in vitro*. Usually, patients were hospitalized one week after the first onset of COVID-19 symptoms. Most frequently patients were hospitalized due to dyspnea. Fifteen patients had a CT-scan showing the specific characteristics of COVID-19 pneumonia. Nine patients did not need oxygen supplementation, but all others received oxygen during their stay. All patients received empiric antimicrobial treatment except four patients. Four patients were treated additionally with oral Oseltamivir until a negative influenza test was available. The median length of hospitalization was 8 (3–108) days. The median age of patients was high (71 years), consistent with the fact that symptomatic COVID-19 disease and hospital admissions are more prevalent in the elderly. Peripheral blood was drawn immediately after hospitalization to analyze T cell responses during acute infection. The laboratory parameters of the patients are depicted in Table 1 (Sup. Tab. 1). All SARS-CoV-2 infections were unequivocally confirmed by certified diagnostic RT-PCRs.

### CD4+ T cells are not cytotoxic during SARS-CoV-2 infection

CD4+ T cells usually function as helper cells, but have been shown to be capable of cytotoxicity after several virus infections, including those with coronaviruses (6, 24). Therefore, we analyzed the production of cytotoxic molecules in CD4+ T cells upon SARS-CoV-2 infection without any additional stimulation of lymphocytes. First, we determined the numbers of CD4+ T cells in the blood of COVID-19 patients and stratified the patients into age groups of 29-79 (median 62) and 80-96 (median 86) years. CD4+ T cell counts were reduced compared to normal clinical references: in the 29-79 age group, the median was 333 CD4+ T cells per μl vs. 555-1460 CD4+ T cells per μl in healthy donors, and in the 80-96 age group, the median was 319 CD4+ T cells per μl vs 540-720 CD4+ T cells per μl in age-matched control individuals (25) (Fig. 1A). No difference in CD4+ T cell counts between the analyzed age groups was observed. Next, we determined the differentiation status of all CD3+CD4+ T cells according to the expression of CD45RO, CCR7, and CD28 and stratified CD4+ T cells into naïve (N, CD45RO− CCR7+ CD28+), central memory (CM, CD45RO+ CCR7+ CD28+), transitional memory (TM, CD45RO+ CCR7− CD28+), effector memory (EM, CD45RO+ CCR7− CD28−), and terminally differentiated effector (E, CD45RO− CCR7− CD28−) subpopulations (Fig. 1B). The gating strategy is shown in Figure S1. Subsequently, we compared the distribution of subpopulations between COVID-19 patients and age-matched healthy controls, again stratified according to age. No obvious differences between COVID-19 patients and healthy controls were found for any of the CD4+ T cell subtypes (Fig. 1C). To characterize their cytotoxic profile, we stained total CD4+ T cells directly *ex vivo* without re-stimulation for the cytotoxic molecules GzmA, GzmB, and perforin and compared the two age groups between COVID-19 patients and age-matched healthy controls (Fig. 1C-F). Again, we did not find clear differences between groups, except that the frequency of GzmB-producing cells was slightly increased in the 29-79 year group of COVID-19 patients compared to healthy controls, yet with largely overlapping confidence intervals (Fig. 1C, E).

**Fig. 1.**
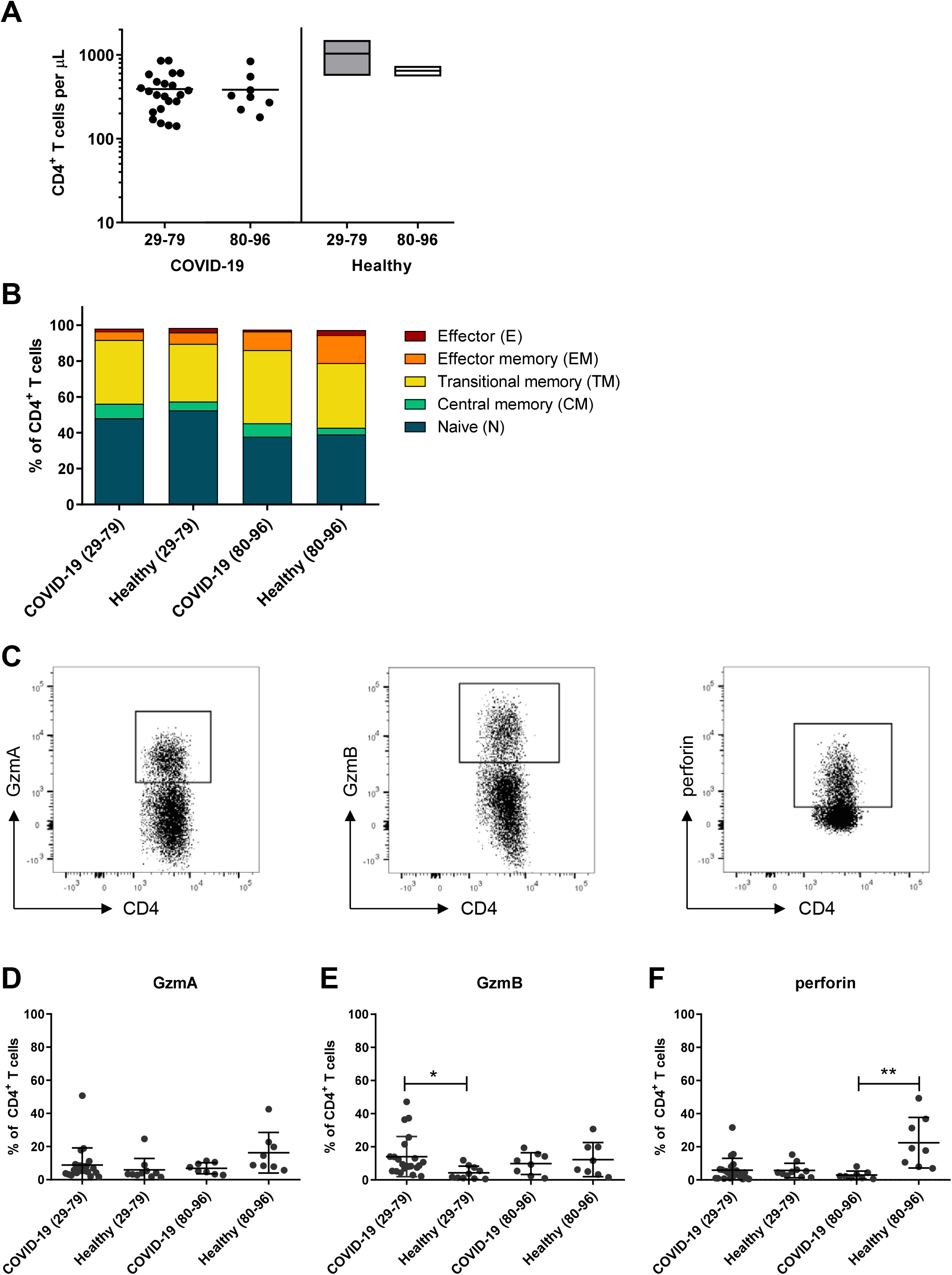
CD4^+^ T cells in COVID-19 patients. CD4^+^ T cells in blood of patients with mild COVID-19 and healthy donors were analyzed by flow cytometry. The concentration of CD4^+^ T cells in peripheral blood of patients at the day of hospitalization and values of healthy donors of different age (**A**). The differentiation status of CD4^+^ T cells was determined by the expression of CD45RO, CCR7, and CD28. CD3^+^ CD4^+^ T cells were divided in naïve (N, CD45RO^−^ CCR7^+^ CD28^+^), central memory (CM, CD45RO^+^ CCR7^+^ CD28^+^), transitional memory (TM, CD45RO^+^ CCR7^−^ CD28^+^), effector memory (EM, CD45RO^+^ CCR7^−^ CD28^−^), and terminally differentiated effector (E, CD45RO^−^ CCR7^−^ CD28^−^) subpopulations (**B**). The production of GzmA, GzmB, and perforin in CD3^+^ CD4^+^ T cells is shown in representative dot plots (**C**). Percentages of CD4^+^ T cells producing GzmA (**D)**, GzmB (**E**), and perforin (**F**). Each dot represents an individual patient. Statistically significant differences are indicated by asterisks (* < 0.05; ** < 0,01; Unpaired t-test with Welch’s correction).

Conversely, perforin responses were reduced in the older age group of COVID-19 patients (Fig. 1C, F). The overall data failed to reveal a meaningful cytotoxic response of CD4+ T cells early after SARS-CoV-2 infection, and we did not further analyze CD4+ cells in the current study.

### Expansion of CD8+ T cells with a cytotoxic profile upon SARS-CoV-2 infection

CD8+ T cell are also named cytotoxic T cells and constitute the main T cell population that can kill virus-infected cells. Therefore, we analyzed the production of cytotoxic molecules in CD8+ T cells in a cohort of SARS-CoV-2-infected individuals.

We first determined CD8+ T cell numbers in the blood of COVID-19 patients in the two age groups. CD8+ T cell counts were clearly reduced in both groups compared to the numbers reported in the literature (25), and an additional significant reduction was found for COVID-19 patients over 80 years of age (Fig. 2A). When calculating the Pearson correlation, we found an inverse association between CD8+ T cell counts in peripheral blood and patient age (Fig. 2B). Next, we determined the distribution of different CD8+ T cell subsets (defined parallel to the criteria for CD4+ T cells described above) and compared COVID-19 patients with age-matched healthy controls, again in two age groups. The gating strategy is shown in Figure S2. Differences were found for the 29-79 year group in which the frequency of naïve CD8+ T cells was clearly reduced in COVID-19 patients, whereas percentages of effector, effector memory, and transitional memory cells were enhanced compared to healthy individuals, suggesting an ongoing CD8+ T cell response in COVID-19 patients (Fig. 2C). Interestingly, this difference was almost absent in the older age group, most likely because the pool of naïve CD8+ T cells largely disappears in elderly individuals (23). To characterize the profile of CD8+ T cells, we stained CD8+ cells for cytotoxic molecules and compared the two age groups from COVID-19 patients and healthy controls (Fig. 3A). Cells were analyzed directly *ex vivo* without any re-stimulation. We found a significant difference between the patients and controls in the younger age group. COVID-19 patients had higher frequencies of CD8+ T cells producing the cytotoxic molecules GzmA and GzmB as well as perforin compared to healthy controls (Fig. 3A-D). Thus, a clear cytotoxic profile could be detected at an early stage of SARS-CoV-2 infection. In the 80-96 age group, an enhanced production of cytotoxic molecules in CD8+ T cells was not evident after infection, most likely because the CD8+ T cells of elderly non-infected individuals already express high background levels of cytotoxic molecules (23). However, if these cells still mediate cytotoxic functions remains elusive.

**Fig. 2.**
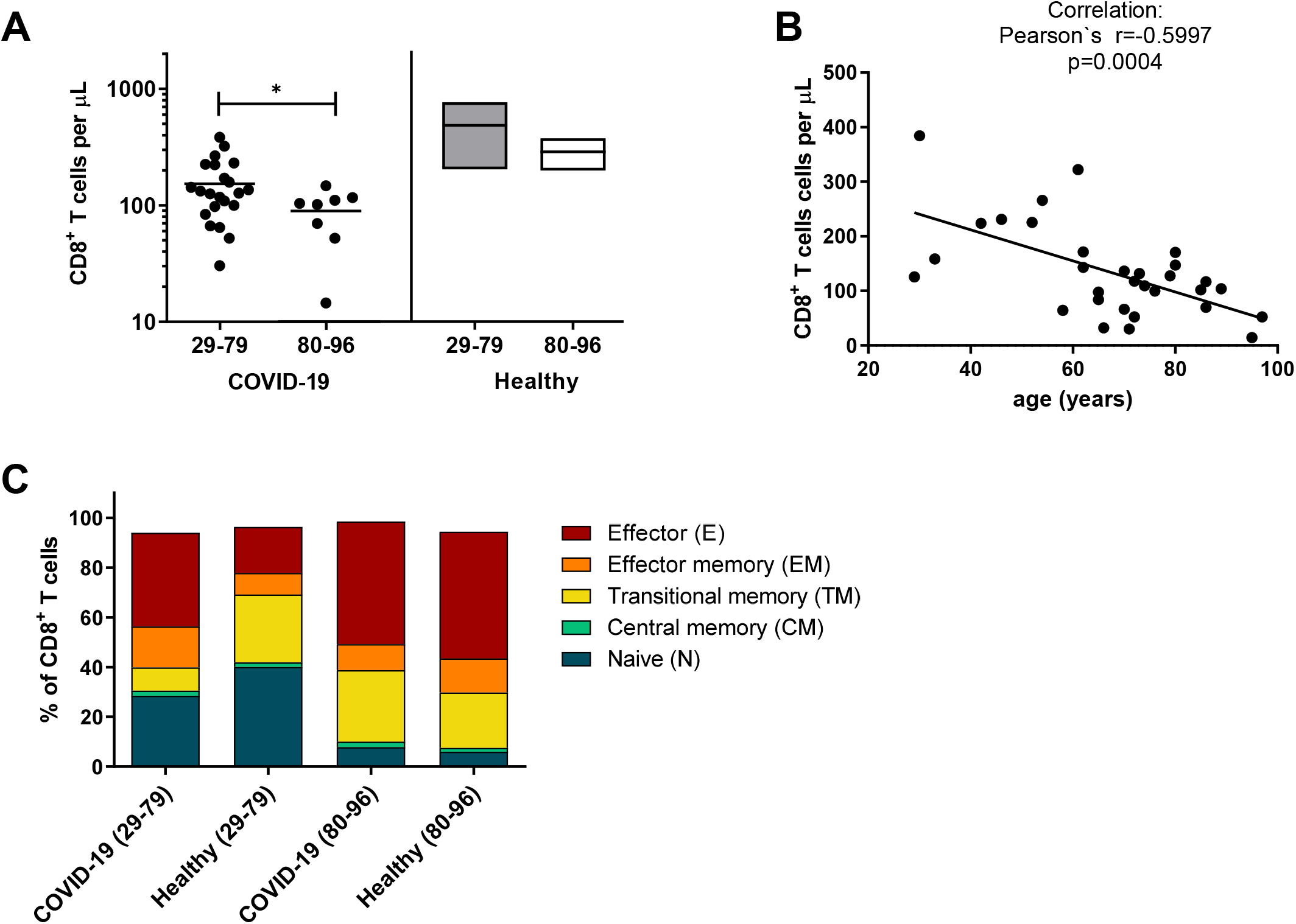
Reduced numbers of circulating CD8^+^ T cells in elderly COVID-19 patients. CD8^+^ T cells in the blood of patients with mild COVID-19 were analyzed by flow cytometry. The concentration of CD8^+^ T cells in peripheral blood of patients at the day of hospitalization and values of healthy donors of different age (**A**). Correlation of age with concentration of CD8^+^ T cells in the blood of acute COVID-18 patients (**B**). Each dot represents an individual patient. The differentiation status of CD8^+^ T cells was determined by the expression of CD45RO, CCR7, and CD28. CD3^+^ CD8^+^ T cells were divided in naïve (N, CD45RO^−^ CCR7^+^ CD28^+^), central memory (CM, CD45RO^+^ CCR7^+^ CD28^+^), transitional memory (TM, CD45RO^+^ CCR7^−^ CD28^+^), effector memory (EM, CD45RO^+^ CCR7^−^ CD28^−^), and terminally differentiated effector (E, CD45RO^−^ CCR7^−^ CD28^−^) subpopulations. Statistically significant differences are indicated by asterisks (* < 0.05; Unpaired t-test with Welch’s correction, Pearson’s correlation coefficient).

**Fig. 3.**
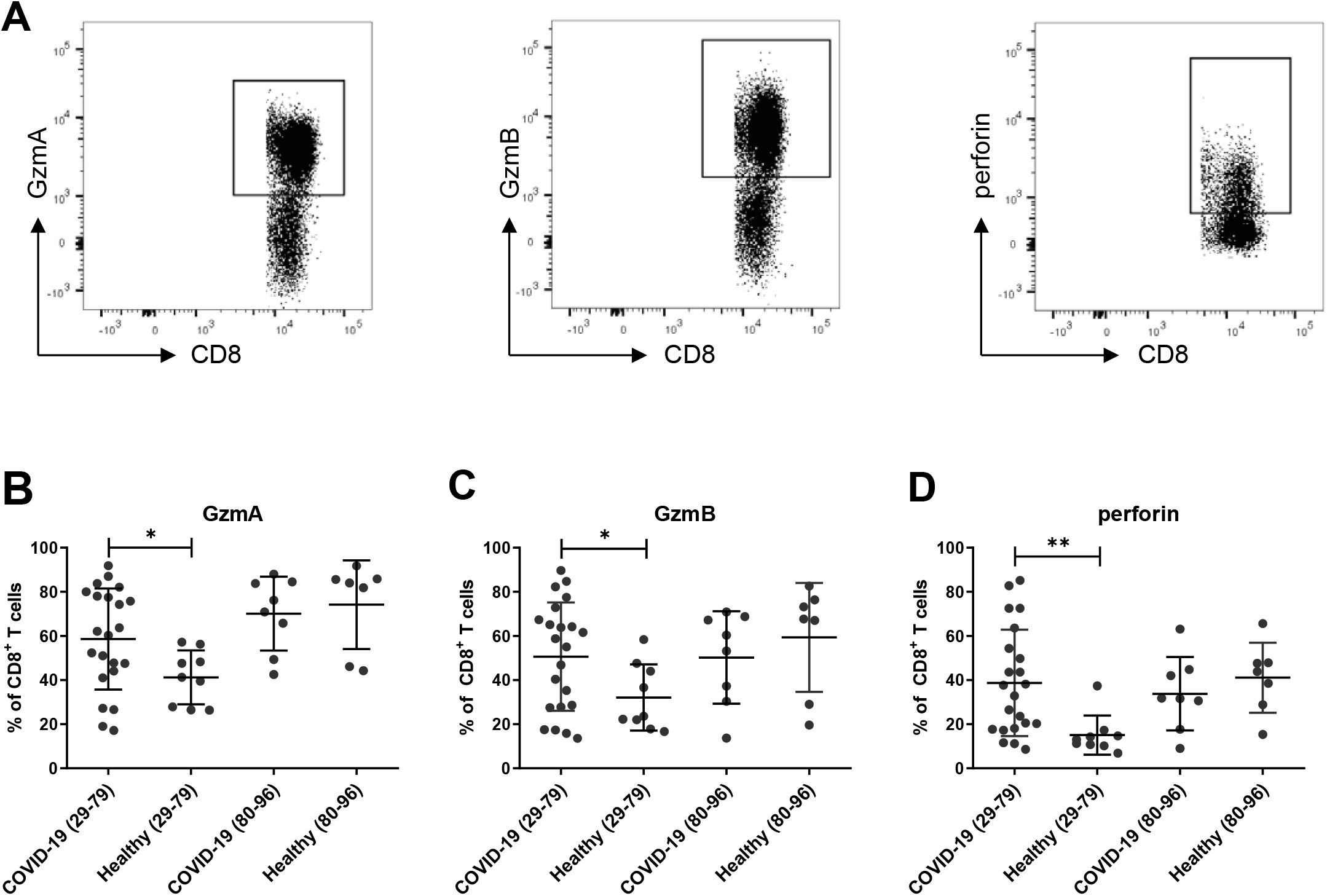
Production of cytotoxic molecules by CD8^+^ T cells. Production of cytotoxic molecules in CD8^+^ T cells in the blood from patients with mild COVID-19 and healthy donors was characterized by flow cytometry. Representative dot plots show the production of GzmA, GzmB, and Perforin in CD8^+^ T cells (**A**). Percentage of CD8^+^ T cells producing GzmA (**B**), GzmB (**C**), and perforin (**D**). Each dot represents an individual patient. Statistically significant differences are indicated by asterisks (* < 0.05; ** < 0.001; Non-parametric Mann-Whitney U test).

Some studies on T cell responses in COVID-19 patients reported that CD8+ T cells may already become functionally exhausted during acute infection (26). This hypothesis was based on the analysis of PD-1 expression by T cells during early COVID-19. In functional terms, PD-1, in conjunction with its ligands PD-L1 and PD-L2, exerts potent immune-inhibitory activities. However, its expression is induced by T cell receptor (TCR) activation (27) and TCR downstream NFAT signaling (28). PD-1 expression is a hallmark of recent TCR-based recognition of MHC-presented antigens that is often up-regulated on cytotoxic effector T cells during acute infections (29, 30). We found that about 20% of total CD8+ T cells expressed PD-1 in healthy controls, as well as in COVID-19 patients (Fig. 4A). Most of these cells expressed GzmA with no apparent differences between the groups (Fig. 4B). However, for GzmB and perforin, we found a higher frequency of positive cells among PD-1+ CD8+ T cells in the group of younger COVID-19 patients compared to healthy controls. This difference was absent for the older age group (Fig. 4C, D). Our data indicate that PD-1+ CD8+ T cells express cytotoxic molecules and should not be misclassified as functionally exhausted T cells during the early SARS-CoV-2 infection.

**Fig. 4.**
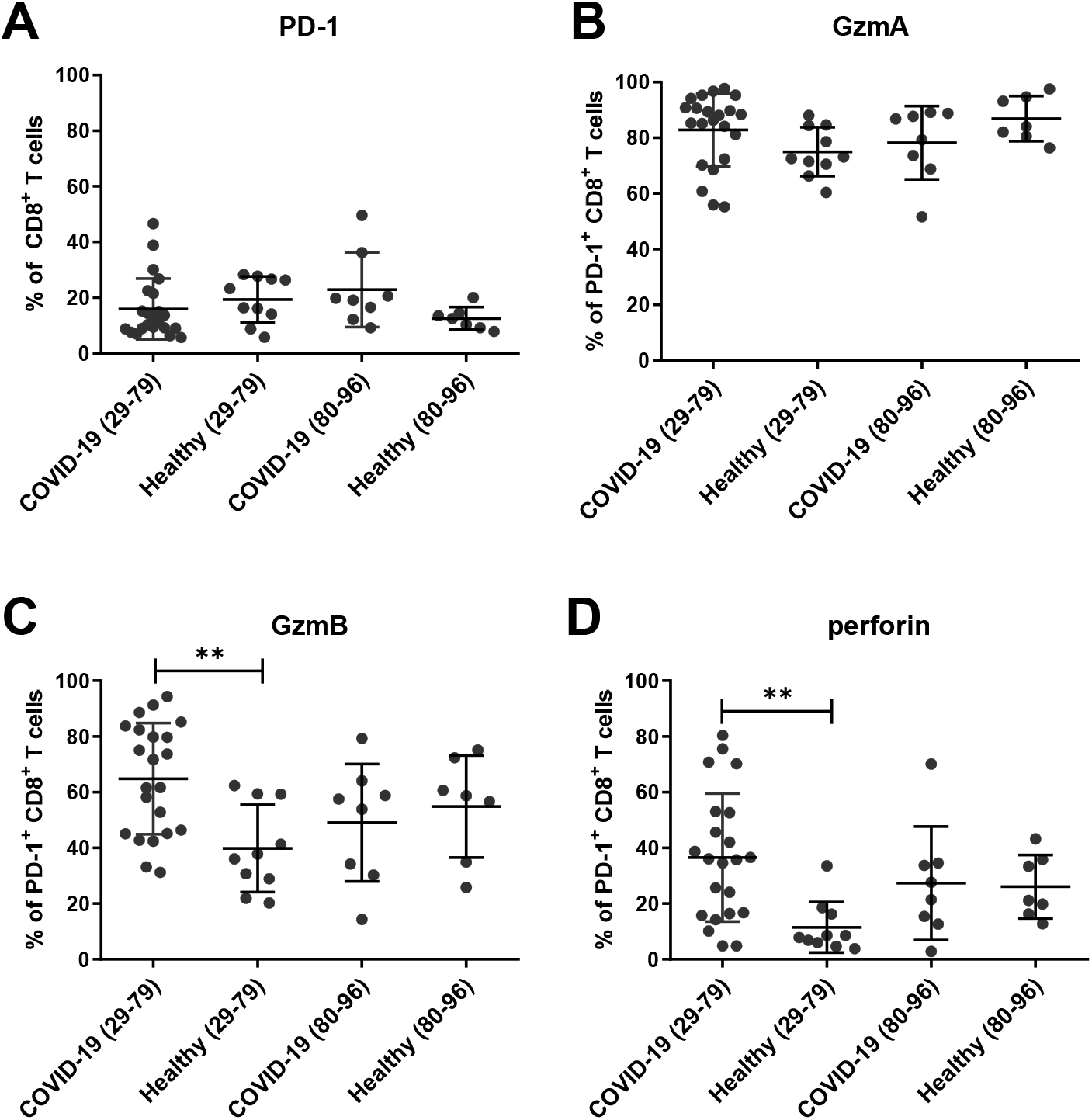
Production of cytotoxic molecules by PD-1^+^ CD8^+^ T cells. Expression of PD-1 and the production of cytotoxic molecules in PD-1^+^ CD8^+^ T cells in the blood of patients with mild COVID-19 was characterized by flow cytometry. Percentages of CD8^+^ PD-1^+^ T cells (**A**) and of PD-1^+^ CD8^+^ T cells producing GzmA (**B**), GzmB (**C**), and perforin (**D**) were calculated. Each dot represents an individual patient. (** < 0.01; Non-parametric Mann-Whitney U test).

Here, we clearly demonstrate a cytotoxic profile in CD8+ T cells upon SARS-CoV-2 infection, which was also found in CD8+ T cells expressing PD-1.

### The frequency of CD8+ T cells with a cytotoxic profile in COVID-19 is reduced with patients’ age

Individual subpopulations of CD8+ T cells differ in their ability to produce cytotoxic molecules, with the highest potency for effector T cell populations. To investigate which CD8+ T cell subpopulation dominates the cytotoxic profile of CD8+ T cells in mild COVID-19 patients, we analyzed the expression of Gzms and perforin in all five T cell subpopulations.

The representative histogram shows that GzmA was produced by transitional memory, effector memory, and effector cells, whereas GzmB and perforin were only found in the latter two populations in our *ex vivo* analysis (Fig. 5A). Next, we assessed whether the production of cytotoxic molecules by effector CD8+ T cell subpopulations is influenced by the age of COVID-19 patients. For a precise analysis of age effects on the expression of cytotoxic molecules we stratified the COVID-19 patients into 3 age groups (29-66 (median 56); 70-76 (median 73); 80-96 years (median 86)). Interestingly, for effector and effector memory cells, the percentages of GzmA-as well as perforin-positive cells were significantly reduced in the 80-96 age group compared to the 29-69 age group (Fig. 5E, G, H, G). For transitional memory cells, this was only the case for GzmA (Fig. 5B). This suggests a functional impairment of the cytotoxic program in the CD8+ T cells of elderly COVID-19 patients.

**Fig. 5.**
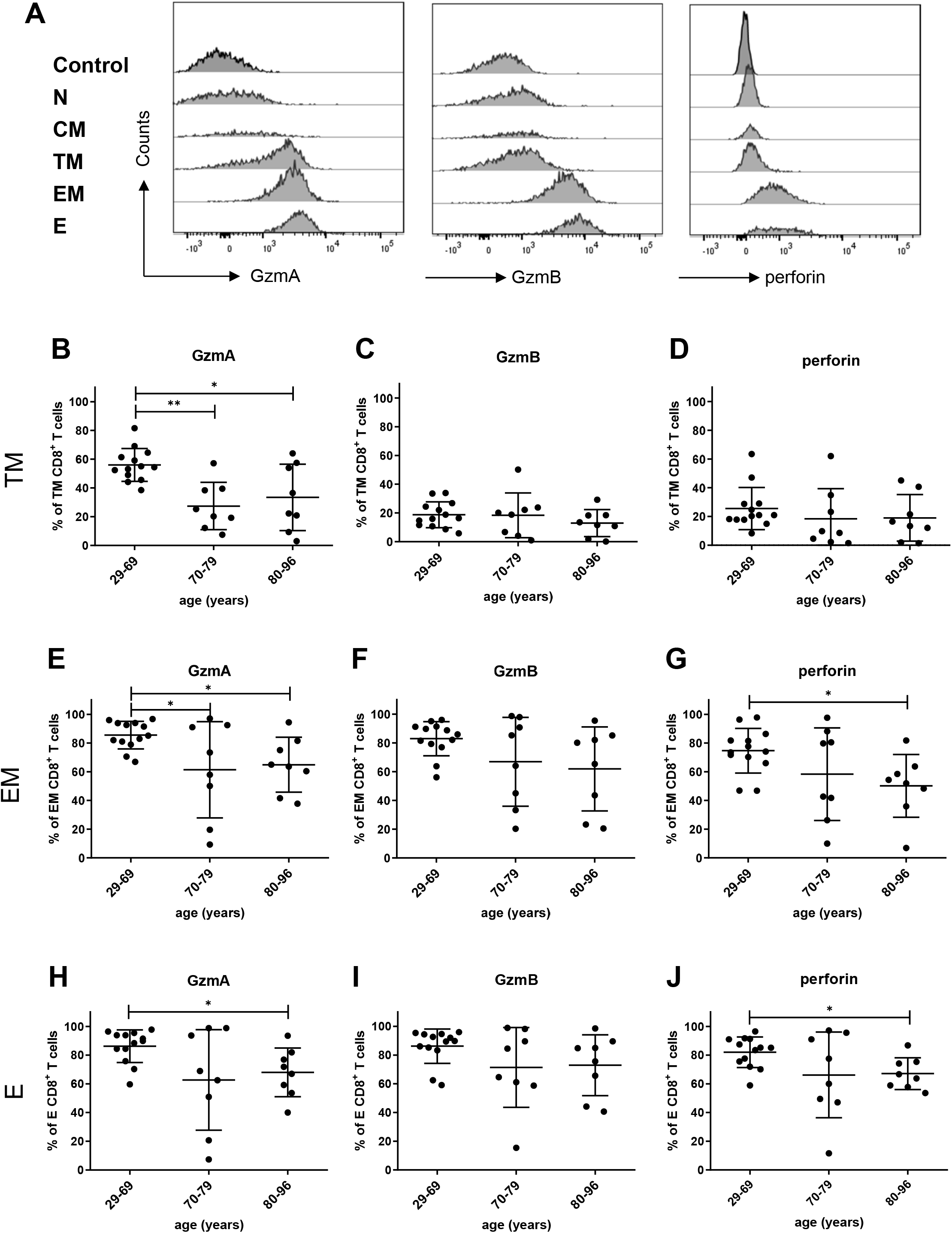
Production of cytotoxic molecules in different subpopulations of CD8^+^ T cells from COVID-19 patients. Differentiation of CD8^+^ T cells in the blood of patients with mild COVID-19 was analyzed by flow cytometry. Representative histograms of the production of GzmA, GzmB, and Perforin in CD8^+^ T cells on different stages of differentiation (**A**). The frequencies of transitional memory (TM, CD45RO^+^ CCR7^−^ CD28^+^), effector memory (EM, CD45RO^+^ CCR7^−^ CD28^−^), and terminally differentiated effector (E, CD45RO^−^ CCR7^−^ CD28^−^) CD8^+^ T cells producing GzmA, GzmB, and perforin in blood of patients with mild COVID-19 disease were detected by flow cytometry. TM CD8^+^ T cells producing GzmA (**B**), GzmB (**C**), and perforin (**D**); EM CD8^+^ T cells producing GzmA (**E**), GzmB (**F**), and perforin (**G**), E CD8^+^ T cells producing GzmA (**H**), GzmB (**I**),and perforin (**J**). Each dot represents an individual patient. Statistically significant differences are indicated by asterisks (* < 0.05; Dunn test with the Benjamini–Hochberg correction for multiple testing).

The simultaneous expression of different cytotoxic molecules is a feature of effector cells with a strong cytolytic potential. Therefore, we also performed single-cell analysis of CD8+ T cells from COVID-19 patients to determine the expression profiles of cytotoxic molecules for the different subpopulations of CD8+ T cells. For transitional memory cells, most cells with a cytotoxic profile produced only GzmA and there was no obvious difference between the age groups (Fig. 6A-C). Surprisingly, the vast majority of effector and effector memory cells produced all three cytotoxic molecules simultaneously (Fig. 6D-I). While all patients from the youngest age group had multifunctional effector cells, some individual patients from the older age groups showed reduced multifunctional responses (Fig. 6E, F, H, I). Interestingly, a comparison of effector CD8+ T cells from age-matched healthy controls and COVID-19 patients revealed that uninfected individuals had two dominating effector cell populations: GzmA and GzmB double-positive as well as GzmA, GzmB and perforin triple-positive cells, whereas in COVID-19 patients, the triple-positive cells unanimously dominated the response (Fig. S3). It remains to be elucidated whether CD8+ T cells that produce multiple cytotoxic molecules are critical for SARS-CoV2 control or the virus-induced immunopathology, or both. However, since our analysis was focused on COVID-19 patients who had a mild disease course without signs of immunopathology, a protective role appears more likely.

**Fig. 6.**
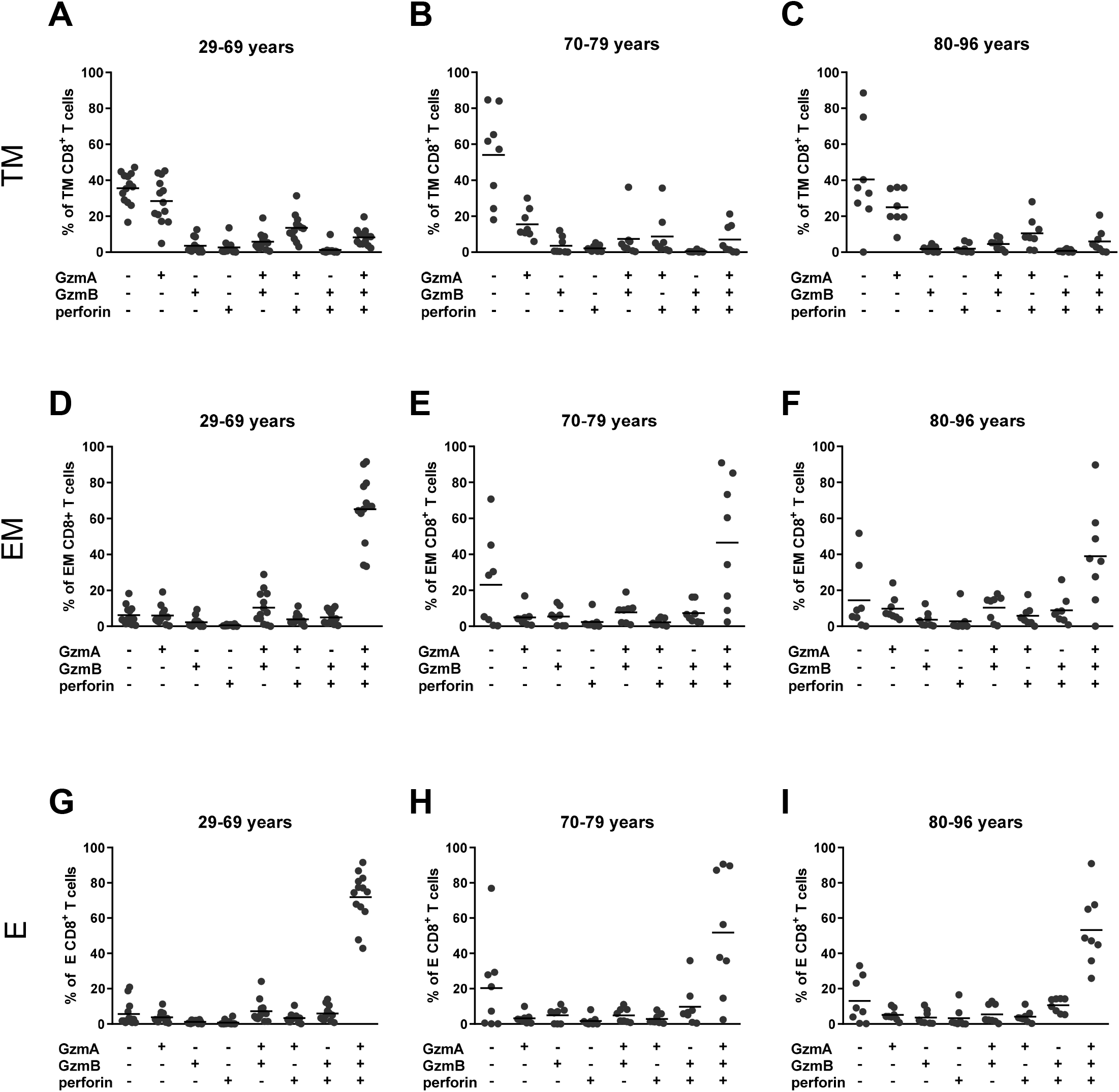
Simultaneous production of GzmA, GzmB, and perforin by CD8^+^ T cells from COVID-19 patients. Differentiation of CD8^+^ T cells in blood of patients with mild COVID-19 was characterized by flow cytometry. The frequencies of CD8^+^ T cells simultaneously producing GzmA, GzmB, and Perforin from patients in the 29-66 years, 70-79 years, 80-96 years age groups were calculated for transitional memory (TM, CD45RO^+^ CCR7^−^ CD28^+^) (**A-C**), effector memory (EM, CD45RO^+^ CCR7^−^ CD28^−^) (**D-F**), and terminally differentiated effector (E, CD45RO^−^ CCR7^−^ CD28^−^) (**G-I**) CD8^+^ T cells. Each dot represents an individual patient.

## Discussion

Cytotoxicity of T cells is the decisive factor for the elimination of virus-infected cells during different acute infections. The CTL-mediated elimination of the virus can also lead to damage of infected organs and to the progression of virus-mediated diseases. A viral infection usually induces a polyclonal activation and expansion of T cells which leads to lymphocytosis in the peripheral blood and a local or systemic lymphadenopathy. Some expanded T cells later differentiate into effector CTLs. In the case of the SARS-CoV-2 infection, the frequency of all lymphocytes in the blood and especially the frequencies of CD8+T cells are strongly reduced in COVID-19 patients (3, 31). In the case of a mild disease course, this reduction was not as pronounced as in the case of severe diseases (31). Interestingly, the recovery during COVID-19 is associated with the reappearance of circulating effector T cells in the blood (32). We analyzed only mild COVID-19 cases here and also found that the frequencies of T cells in the blood were not greatly reduced, except in the older patient group. Our *ex vivo* study indicates that COVID-19 patients show a cytotoxic response dominated by CD8+ T cell in contrast to CD4+ T cells during the early period of infection. It is very likely that this CD8+ T cell response contributes to virus control and its subsequent elimination as it has been shown in other viral infections affecting the respiratory tract (33–35). However, no cytotoxic CD4+ T cell response could be demonstrated during SARS-CoV-2 infection, which may not be too surprising since these cells were mainly found in chronic and not in acute self-limiting viral infections (36). Interestingly, PD-1-positive CD8+ T cells show a clear profile of cytotoxicity, indicating that they are most likely not functionally exhausted during acute SARS-CoV-2 infection. This is in clear contrast to suggestions by other groups (26) but in line with previous findings made by us and others indicating that PD-1 is a marker for activation rather than for exhaustion during early phases of infections (30) and is especially up-regulated on T cell subsets that produce cytotoxic molecules such as Gzms and perforin. PD-1 expression is induced upon T cell activation and TCR signaling and only sets the stage for subsequent immune checkpoint control post-acute infection. This is then strongly influenced by the presence of the PD-1 ligands on virus-infected target cells (21). The control of cytotoxic cells is necessary during later phases of the immune response, where immunopathology rather than viral replication may become the greatest danger.

In contrast to the younger age group, no clear cytotoxic CD8+ T cell response in the blood could be demonstrated in the age group of over 80 year old patients. Two main points influence this finding: the age-dependent reduction of CD8+ T cell frequencies in the blood of COVID-19 patients (25) and the previously reported high baseline expression levels of Gzms and perforin being a feature of senescent CD8+ T cells from elderly individuals (37).

For an aged immune system, the reduction of T lymphocytes and processes of immune senescence are characteristic (37). However, the nature of the progressive loss of circulating CD8+ T cells in elderly COVID-19 patients is not completely understood. One possible explanation may be an enhanced migration of T cells from the blood into the infected tissue. Usually, the accumulation of T lymphocytes leads to a progressive inflammation in the infected organs. SARS-CoV-2 infects lung epithelial cells, which might recruit cytotoxic T cells into the lung. In the early phase of infection, which we analyzed here, they most like contribute to virus control in the lung. However, sustained T cell cytotoxicity might also contribute to organ damage. Thus, the precise recognition and elimination of infected cells without the induction of too much inflammation and tissue destruction is necessary for the survival of infected patients. This delicate balance of two opposing processes is very important for survival. Here, multifunctional T cells, producing both perforin and Gzms at the same time may be very important, as it has been shown that perforin is a critical enabler of the apoptotic effects mediated by Gzms. Cells producing perforin and Gzms are necessary for the efficient control of virus infections (20). We found many T cells producing Gzms and perforin in our COVID-19 patient cohort, although their frequencies were reduced in elderly patients. They might have contributed to efficient virus control, since all our patients showed only mild symptoms and fully recovered from COVID-19. Multifunctional cytotoxic T cells often express PD-1 (38), not because they are functionally impaired during acute infection, but as an important negative switch to shut them down when responses are either too strong or maintained for too long.

The cytotoxic molecules analyzed here share some overlapping functions but also elicit non-redundant features. The critical effector molecule for target cell killing is perforin, as it promotes the entry of Gzms into target cells and in this way enables the cytotoxic functionality of Gzms (20). Thus, CTLs producing Gzms without perforin can induce severe inflammation triggered by the aimless release of Gzms and inflammatory cytokines. Once Gzms enter infected target cells in the presence of perforin, they mediate apoptosis of these cells. Thus, it is tempting to speculate that the herein identified age-associated reduction of CTLs expressing perforin may be an additional factor in COVID-19 progression, as it might support lung inflammation.

Our current data support the concept that cytotoxic CD8+ T cells play an important role in the control of early SARS-CoV-2 infections, but may also be a factor of immune pathogenesis and COVID-19 progression during later periods of infection. Thus, it will be important to carefully balance therapeutic measures either supporting or suppressing T cell responses in future COVID-19 therapy. Recent suggestions to therapeutically administer checkpoint inhibitors, which are efficiently used for tumor immune therapy, for the treatment of COVID-19 patients (39) should be reevaluated, since we did not find functionally exhausted CD8+ T cells in our patients. In agreement with our previous findings based on acute virus infection models in mice (30, 38), our clinical study suggests that a checkpoint therapy might enhance the functionality of the PD-1-expressing cytotoxic CD8+ T cells in COVID-19 patients and improve virus control, but with a potential to exaggerate the immunopathology in the lung and other organs, which might actually accelerate decompensation.

## Material and Methods

### Study population and design

For this study, 30 patients with mild COVID-19 cases were recruited directly after hospitalization, which occurred approximately one week after symptom onset and at least one positive SARS-CoV-2 PCR result was available (SARS-CoV-2 test, Altona Diagnostics, Hamburg). Written consent was obtained from each of the study participants. The study was approved by the University Hospital Essen’s ethical committee (ethics vote 20-9216-BO). Clinical characteristics of all patients are shown in Table 1. Two groups of age-matched healthy individuals were used as controls: 10 individuals (6F/4M), median age 50.3 years, median BMI 24.3, no diabetes and 8 individuals (3F/5M), median age 85.2 years, median BMI 25.0, including 3 with diabetes. Otherwise no medical conditions were reported for the control group.

### Preparation of PBMCs

Peripheral blood mononuclear cells (PBMCs) were isolated from peripheral blood by gradient centrifugation. Blood was collected in EDTA S-monovettes (Sarstedt). Collected blood was pre-diluted with RPMI-1640 (Thermo Fisher Scientific) and overlaid onto 15 mL Pancoll separating solution (PAN-Biotech). The tubes were centrifuged at 1600 rcf for 15 min at room temperature with deceleration of the centrifuge set to low. Isolated PBMCs were washed twice with RPMI-1640.

### Cell surface and intracellular staining by flow cytometry

Surface and intracellular staining were performed as described previously (40). For the surface staining of human cells, specific antibodies against human CD3 (OKT3, BioLegend), CD4 (OKT4, BioLegend), CD8 (BW135/80, Miltenyi Biotec), CD45RO (UCHL1, BioLegend), CCR7 (G043H7, BioLegend), CD28 (CD28.2, BioLegend), and PD-1 (EH12.2 H7, BioLegend) were used. For intracellular staining antibodies against human GzmA (CB9, BioLegend), GzmB (QA16A02, BioLegend) and perforin (B-D48, BioLegend) were used. Dead cells were determined by Fixable Viability Dye (Thermo Fisher) staining and excluded from analysis. The numbers of CD3+CD4+ and CD3+CD8+ T cells in the blood were calculated from lymphocytes counts measured in a certified clinical laboratory for every patient.

Data were acquired on a LSR II flow cytometer (Becton Dickinson) from 250,000-300,000 lymphocyte-gated events per sample. Analyses were done using FACSDiva software (Becton Dickinson) and FlowJo software (Becton Dickinson).

### Statistical Analysis

Statistics comparing two groups were done using the unpaired non-parametric t-test or Mann-Whitney U test. A Pearson correlation coefficient was used for the definition of correlation. (GraphPad Prism software; GraphPad Software Inc., San Diego, USA). When more than two groups were compared, a Dunn test with the Benjamini–Hochberg correction for multiple testing was performed (R-package dunn.test, version 1.3.4).

## Acknowledgements

We are grateful to the Stiftung Universitätsmedizin Essen of University Hospital Essen for the financial support of this study and we are thankful to Westdeutsche Biobank Essen for material sampling.

---

**Supplement Table 1. Laboratory characteristics of patients according to age.**

30 patients with mild COVID-19 cases were recruited directly after hospitalization, which occurred approximately one week after symptom onset and at least one positive SARS-CoV-2 PCR result was available. Patients were stratified into two age groups of 29-79 and 80-96 years.

**Supplement Fig. S1. Gating strategy for CD4^+^ T cell subpopulations.**

Exemplary gating strategy for definition of CD4^+^ T cell subpopulations. Naïve (N, CCR7^+^ CD45RO^−^ CD28^+^), central memory (CM, CCR7^−^ CD45RO^+^ CD28^+^), transitional memory (TM, CCR7^−^ CD45RO^+^ CD28^+^), effector memory (EM, CCR7^−^ CD45RO^+^ CD28^−^), and effector (E, CCR7-CD45RO^−^ CD28^−^) CD4^+^ T cell subpopulations were characterized using CCR7, CD45RO, and CD28.

**Supplement Fig. S2. Gating strategy for CD8^+^ T cell subpopulations and simultaneous production of cytotoxic molecules.**

Exemplary gating strategy for definition of CD8^+^ T cell subpopulations and the simultaneous production of GzmA, GzmB and perforin. Naïve (N, CCR7^+^ CD45RO^−^ CD28^+^), central memory (CM, CCR7^+^ CD45RO^+^ CD28^+^), transitional memory (TM, CCR7^−^ CD45RO^+^ CD28^+^), effector memory (EM, CCR7^−^ CD45RO^+^ CD28^−^), and effector (E, CCR7-CD45RO^−^ CD28^−^) CD8^+^ T cell subpopulations were characterized using CCR7, CD45RO, and CD28.

**Supplement Fig. S3. Simultaneous production of GzmA, GzmB, and Perforin in CD8^+^ T cells from COVID-19 patients and healthy controls.**

The simultaneous production of GzmA, GzmB, and perforin by CD8^+^ T cells in blood of patients with mild COVID-19 and healthy controls was characterized by flow cytometry. The frequencies of CD8^+^ T cells producing GzmA, GzmB, and perforin from patients in the 29-79 years and 80-96 years age groups were calculated for effector memory (EM, CD45RO^+^ CCR7^−^ CD28^−^) (**A, B**), and terminally differentiated effector (E, CD45RO^−^ CCR7^−^ CD28^−^) (**C, D**) CD8^+^ T cells. Statistically significant differences are indicated by asterisks (* < 0.05; ** < 0.01; *** < 0.001; Non-parametric Mann-Whitney U test).

## References

1. Docherty AB, Harrison EM, Green CA, Hardwick HE, Pius R, Norman L, Holden KA, Read JM, Dondelinger F, Carson G, Merson L, Lee J, Plotkin D, Sigfrid L, Halpin S, Jackson C, Gamble C, Horby PW, Nguyen-Van-Tam JS, Ho A, Russell CD, Dunning J, Openshaw PJ, Baillie JK, Semple MG. 2020. Features of 20 133 UK patients in hospital with covid-19 using the ISARIC WHO Clinical Characterisation Protocol: prospective observational cohort study. BMJ 369:m1985.

2. Guan WJ, Ni ZY, Hu Y, Liang WH, Ou CQ, He JX, Liu L, Shan H, Lei CL, Hui DSC, Du B, Li LJ, Zeng G, Yuen KY, Chen RC, Tang CL, Wang T, Chen PY, Xiang J, Li SY, Wang JL, Liang ZJ, Peng YX, Wei L, Liu Y, Hu YH, Peng P, Wang JM, Liu JY, Chen Z, Li G, Zheng ZJ, Qiu SQ, Luo J, Ye CJ, Zhu SY, Zhong NS, China Medical Treatment Expert Group for C. 2020. Clinical Characteristics of Coronavirus Disease 2019 in China. N Engl J Med 382:1708–1720.

3. Liu K, Chen Y, Lin R, Han K. 2020. Clinical features of COVID-19 in elderly patients: A comparison with young and middle-aged patients—. J Infect 80:e14–e18.

4. Verity R, Okell LC, Dorigatti I, Winskill P, Whittaker C, Imai N, Cuomo-Dannenburg G, Thompson H, Walker PGT, Fu H, Dighe A, Griffin JT, Baguelin M, Bhatia S, Boonyasiri A, Cori A, Cucunuba Z, FitzJohn R, Gaythorpe K, Green W, Hamlet A, Hinsley W, Laydon D, Nedjati-Gilani G, Riley S, van Elsland S, Volz E, Wang H, Wang Y, Xi X, Donnelly CA, Ghani AC, Ferguson NM. 2020. Estimates of the severity of coronavirus disease 2019: a model-based analysis. Lancet Infect Dis 20:669–677.

5. Vabret N, Britton GJ, Gruber C, Hegde S, Kim J, Kuksin M, Levantovsky R, Malle L, Moreira A, Park MD, Pia L, Risson E, Saffern M, Salome B, Esai Selvan M, Spindler MP, Tan J, van der Heide V, Gregory JK, Alexandropoulos K, Bhardwaj N, Brown BD, Greenbaum B, Gumus ZH, Homann D, Horowitz A, Kamphorst AO, Curotto de Lafaille MA, Mehandru S, Merad M, Samstein RM, Sinai Immunology Review P. 2020. Immunology of COVID-19: Current State of the Science. Immunity 52:910–941.

6. Malyshkina A, Littwitz-Salomon E, Sutter K, Zelinskyy G, Windmann S, Schimmer S, Paschen A, Streeck H, Hasenkrug KJ, Dittmer U. 2017. Fas Ligand-mediated cytotoxicity of CD4+ T cells during chronic retrovirus infection. Sci Rep 7:7785.

7. Wiesel M, Walton S, Richter K, Oxenius A. 2009. Virus-specific CD8 T cells: activation, differentiation and memory formation. APMIS 117:356–381.

8. Schmidt ME, Varga SM. 2018. The CD8 T Cell Response to Respiratory Virus Infections. Frontiers in Immunology 9;9:678.

9. Chen G, Wu D, Guo W, Cao Y, Huang D, Wang H, Wang T, Zhang X, Chen H, Yu H, Zhang X, Zhang M, Wu S, Song J, Chen T, Han M, Li S, Luo X, Zhao J, Ning Q. 2020. Clinical and immunological features of severe and moderate coronavirus disease 2019. J Clin Invest 130:2620–2629.

10. Nie S, Zhao X, Zhao K, Zhang Z, Zhang Z, Zhang Z. 2020. Metabolic disturbances and inflammatory dysfunction predict severity of coronavirus disease 2019 (COVID-19): a retrospective study. medRxiv doi:10.1101/2020.03.24.20042283:2020.03.24.20042283.

11. Liao M, Liu Y, Yuan J, Wen Y, Xu G, Zhao J, Cheng L, Li J, Wang X, Wang F, Liu L, Amit I, Zhang S, Zhang Z. 2020. Single-cell landscape of bronchoalveolar immune cells in patients with COVID-19. Nature Medicine 26:842–844.

12. Xu Z, Shi L, Wang Y, Zhang J, Huang L, Zhang C, Liu S, Zhao P, Liu H, Zhu L, Tai Y, Bai C, Gao T, Song J, Xia P, Dong J, Zhao J, Wang F-S. 2020. Pathological findings of COVID-19 associated with acute respiratory distress syndrome. The Lancet Respiratory Medicine 8:420–422.

13. Anft M, Paniskaki K, Blazquez-Navarro A, Doevelaar AAN, Seibert F, Hoelzer B, Skrzypczyk S, Kohut E, Kurek J, Zapka J, Wehler P, Kaliszczyk S, Bajda S, Thieme C, Roch T, Konik MJ, Brenner T, Tempfer C, Watzl C, Dolff S, Dittmer U, Westhoff T, Witzke O, Stervbo U, Babel N. 2020. COVID-19 progression is potentially driven by T cell immunopathogenesis. medRxiv doi:10.1101/2020.04.28.20083089:2020.04.28.20083089.

14. Ni L, Ye F, Cheng ML, Feng Y, Deng YQ, Zhao H, Wei P, Ge J, Gou M, Li X, Sun L, Cao T, Wang P, Zhou C, Zhang R, Liang P, Guo H, Wang X, Qin CF, Chen F, Dong C. 2020. Detection of SARS-CoV-2-Specific Humoral and Cellular Immunity in COVID-19 Convalescent Individuals. Immunity 52:971–977 e3.

15. Thieme CJ, Anft M, Paniskaki K, Blazquez-Navarro A, Doevelaar A, Seibert FS, Hoelzer B, Konik MJ, Brenner T, Tempfer C, Watzl C, Dolff S, Dittmer U, Westhoff TH, Witzke O, Stervbo U, Roch T, Babel N. 2020. The SARS-CoV-2 T-cell immunity is directed against the spike, membrane, and nucleocapsid protein and associated with COVID 19 severity. medRxiv doi:10.1101/2020.05.13.20100636:2020.05.13.20100636.

16. Weiskopf D, Schmitz KS, Raadsen MP, Grifoni A, Okba NMA, Endeman H, van den Akker JPC, Molenkamp R, Koopmans MPG, van Gorp ECM, Haagmans BL, de Swart RL, Sette A, de Vries RD. 2020. Phenotype and kinetics of SARS-CoV-2-specific T cells in COVID-19 patients with acute respiratory distress syndrome. Sci Immunol 5.

17. Grifoni A, Weiskopf D, Ramirez SI, Mateus J, Dan JM, Moderbacher CR, Rawlings SA, Sutherland A, Premkumar L, Jadi RS, Marrama D, de Silva AM, Frazier A, Carlin AF, Greenbaum JA, Peters B, Krammer F, Smith DM, Crotty S, Sette A. 2020. Targets of T Cell Responses to SARS-CoV-2 Coronavirus in Humans with COVID-19 Disease and Unexposed Individuals. Cell 181:1489–1501.e15.

18. Le Bert N, Tan AT, Kunasegaran K, Tham CYL, Hafezi M, Chia A, Chng MHY, Lin M, Tan N, Linster M, Chia WN, Chen MI, Wang LF, Ooi EE, Kalimuddin S, Tambyah PA, Low JG, Tan YJ, Bertoletti A. 2020. SARS-CoV-2-specific T cell immunity in cases of COVID-19 and SARS, and uninfected controls. Nature doi:10.1038/s41586-020-2550-z.

19. Snow AL, Pandiyan P, Zheng L, Krummey SM, Lenardo MJ. 2010. The power and the promise of restimulation-induced cell death in human immune diseases. Immunological Reviews 236:68–82.

20. Voskoboinik I, Whisstock JC, Trapani JA. 2015. Perforin and granzymes: function, dysfunction and human pathology. Nature Reviews Immunology 15:388–400.

21. David P, Drabczyk-Pluta M, Pastille E, Knuschke T, Werner T, Honke N, Megger DA, Akhmetzyanova I, Shaabani N, Eyking-Singer A, Cario E, Kershaw O, Gruber AD, Tenbusch M, Dietze KK, Trilling M, Liu J, Schadendorf D, Streeck H, Lang KS, Xie Y, Zimmer L, Sitek B, Paschen A, Westendorf AM, Dittmer U, Zelinskyy G. 2020. Combination immunotherapy with anti-PD-L1 antibody and depletion of regulatory T cells during acute viral infections results in improved virus control but lethal immunopathology. PLoS Pathog 16:e1008340.

22. Xu Z, Shi L, Wang Y, Zhang J, Huang L, Zhang C, Liu S, Zhao P, Liu H, Zhu L, Tai Y, Bai C, Gao T, Song J, Xia P, Dong J, Zhao J, Wang FS. 2020. Pathological findings of COVID-19 associated with acute respiratory distress syndrome. Lancet Respir Med 8:420–422.

23. Vicente R, Mausset-Bonnefont A-L, Jorgensen C, Louis-Plence P, Brondello J-M. 2016. Cellular senescence impact on immune cell fate and function. Aging Cell 15:400–406.

24. Phetsouphanh C, Pillai S, Zaunders JJ. 2017. Editorial: Cytotoxic CD4+ T Cells in Viral Infections. Frontiers in Immunology 4;8:1729.

25. Huppert F, Solomou W, O’Connor S, Morgan K, Sussams P, Brayne C. 1998. Aging and lymphocyte subpopulations: whole-blood analysis of immune markers in a large population sample of healthy elderly individuals. Experimental Gerontology 33:593–600.

26. Diao B, Wang C, Tan Y, Chen X, Liu Y, Ning L, Chen L, Li M, Liu Y, Wang G, Yuan Z, Feng Z, Zhang Y, Wu Y, Chen Y. 2020. Reduction and Functional Exhaustion of T Cells in Patients With Coronavirus Disease 2019 (COVID-19). Front Immunol 11:827.

27. Chikuma S, Terawaki S, Hayashi T, Nabeshima R, Yoshida T, Shibayama S, Okazaki T, Honjo T. 2009. PD-1-mediated suppression of IL-2 production induces CD8+ T cell anergy in vivo. J Immunol 182:6682–9.

28. Simon S, Labarriere N. 2018. PD-1 expression on tumor-specific T cells: Friend or foe for immunotherapy? OncoImmunology 7:e1364828.

29. Jubel JM, Barbati ZR, Burger C, Wirtz DC, Schildberg FA. 2020. The Role of PD-1 in Acute and Chronic Infection. Frontiers in Immunology 24;11:487.

30. Zelinskyy G, Myers L, Dietze KK, Gibbert K, Roggendorf M, Liu J, Lu M, Kraft AR, Teichgraber V, Hasenkrug KJ, Dittmer U. 2011. Virus-specific CD8+ T cells upregulate programmed death-1 expression during acute friend retrovirus infection but are highly cytotoxic and control virus replication. J Immunol 187:3730–7.

31. Liu J, Li S, Liu J, Liang B, Wang X, Wang H, Li W, Tong Q, Yi J, Zhao L, Xiong L, Guo C, Tian J, Luo J, Yao J, Pang R, Shen H, Peng C, Liu T, Zhang Q, Wu J, Xu L, Lu S, Wang B, Weng Z, Han C, Zhu H, Zhou R, Zhou H, Chen X, Ye P, Zhu B, Wang L, Zhou W, He S, He Y, Jie S, Wei P, Zhang J, Lu Y, Wang W, Zhang L, Li L, Zhou F, Wang J, Dittmer U, Lu M, Hu Y, Yang D, Zheng X. 2020. Longitudinal characteristics of lymphocyte responses and cytokine profiles in the peripheral blood of SARS-CoV-2 infected patients. EBioMedicine 55:102763.

32. Odak I, Barros-Martins J, Bošnjak B, Stahl K, David S, Wiesner O, Busch M, Hoeper MM, Pink I, Welte T, Cornberg M, Stoll M, Goudeva L, Blasczyk R, Ganser A, Prinz I, Förster R, Koenecke C, Schultze-Florey CR. 2020. Reappearance of effector T cells is associated with recovery from COVID-19. EBioMedicine 57:102885.

33. Melendi GA, Zavala F, Buchholz UJ, Boivin G, Collins PL, Kleeberger SR, Polack FP. 2007. Mapping and characterization of the primary and anamnestic H-2(d)-restricted cytotoxic T-lymphocyte response in mice against human metapneumovirus. J Virol 81:11461–7.

34. Topham DJ, Tripp RA, Doherty PC. 1997. CD8+ T cells clear influenza virus by perforin or Fas-dependent processes. J Immunol 159:5197–200.

35. Schmidt ME, Varga SM. 2018. The CD8 T Cell Response to Respiratory Virus Infections. Front Immunol 9:678.

36. Hasenkrug KJ, Chougnet CA, Dittmer U. 2018. Regulatory T cells in retroviral infections. PLoS Pathog 14:e1006776.

37. Xu W, Larbi A. 2017. Markers of T Cell Senescence in Humans. Int J Mol Sci 18.

38. David P, Megger DA, Kaiser T, Werner T, Liu J, Chen L, Sitek B, Dittmer U, Zelinskyy G. 2019. The PD-1/PD-L1 Pathway Affects the Expansion and Function of Cytotoxic CD8(+) T Cells During an Acute Retroviral Infection. Front Immunol 10:54.

39. Di Cosimo S, Malfettone A, Pérez-García JM, Llombart-Cussac A, Miceli R, Curigliano G, Cortés J. 2020. Immune checkpoint inhibitors: a physiology-driven approach to the treatment of coronavirus disease 2019. European journal of cancer (Oxford, England : 1990) 135:62–65.

40. Dietze KK, Zelinskyy G, Gibbert K, Schimmer S, Francois S, Myers L, Sparwasser T, Hasenkrug KJ, Dittmer U. 2011. Transient depletion of regulatory T cells in transgenic mice reactivates virus-specific CD8+ T cells and reduces chronic retroviral set points. Proc Natl Acad Sci U S A 108:2420–5.

